# Comparative genomics unravels mechanisms of genetic adaptation for the catabolism of the phenylurea herbicide linuron in *Variovorax*

**DOI:** 10.1101/759100

**Authors:** Başak Öztürk, Johannes Werner, Jan P. Meier-Kolthoff, Boyke Bunk, Cathrin Spröer, Dirk Springael

## Abstract

Biodegradation of the phenylurea herbicide linuron appears a specialization within a specific clade of the *Variovorax* genus. The linuron catabolic ability is likely acquired by horizontal gene transfer but the mechanisms involved are not known. The full genome sequences of six linuron degrading *Variovorax* strains isolated from geographically distant locations were analyzed to acquire insight in the mechanisms of genetic adaptation towards linuron metabolism in *Variovorax*. Whole genome sequence analysis confirmed the phylogenetic position of the linuron degraders in a separate clade within *Variovorax* and indicated their unlikely origin from a common ancestral linuron degrader. The linuron degraders differentiated from non-degraders by the presence of multiple plasmids of 20 to 839 kb, including plasmids of unknown plasmid groups. The linuron catabolic gene clusters showed (i) high conservation and synteny and (ii) strain-dependent distribution among the different plasmids. All were bordered by IS*1071* elements forming composite transposon structures appointing IS*1071* as key for catabolic gene recruitment. Most of the strain carried at least one broad host range plasmid that might have been a second instrument for catabolic gene acquisition. We conclude that clade 1*Variovorax* strains, despite their different geographical origin, made use of a limited genetic repertoire to acquire linuron biodegradation.

**Importance:** The genus *Variovorax* and especially a clade of strains that phylogenetically separates from the majority of *Variovorax* species, appears to be a specialist in the biodegradation of the phenyl urea herbicide linuron. Horizontal gene transfer (HGT) likely played an essential role in the genetic adaptation of those strain to acquire the linuron catabolic genotype. However, we do not know the genetic repertoire involved in this adaptation both regarding catabolic gene functions as well as gene functions that promote HGT neither do we know how this varies between the different strains. These questions are addressed in this paper by analyzing the full genome sequences of six linuron degrading *Variovorax* strains. This knowledge is important for understanding the mechanisms that steer world-wide genetic adaptation in a particular species and this for a particular phenotypic trait as linuron biodegradation.

## Introduction

Linuron [3-(3,4-dichlorophenyl)-1-methoxy-1-methyl urea] is a phenylurea herbicide that has been widely used for weed control in agriculture. Biodegradation is the major route of linuron dissipation in the environment(1). Bacteria belonging to the genus *Variovorax* were isolated from geographically-distant locations either as single strains (2–4) or as members of consortia (4, 5) that have the ability to mineralize and utilize the herbicide for growth. Single strains convert linuron to CO_2_ and cell material while in consortia, *Variovorax* perform particularly the initial hydrolysis of linuron into the primary metabolite 3,4-dichloroaniline (DCA). The metabolic pathway of linuron degradation in *Variovorax* sp. WDL1 and SRS16 are well studied. The linuron hydrolases HylA (identified in WDL1) (6) and LibA (identified in SRS16) (1) perform the hydrolysis of linuron into DCA and *N,O*-dimethylhydroxylamine (*N,O*-DMHA) (1, 6). In both strains, a multicomponent chloroaniline dioxygenase DcaQTA1A2B converts DCA to 4,5-dichlorocatechol while chlorocatechol is further metabolized to oxo-adipate by enzymes encoded by the *ccdCFDE* gene cluster(7). PCR analysis has shown that other linuron-degrading *Variovorax* share the same catabolic genes. Interestingly, based on 16S rRNA gene phylogeny, the linuron degrading *Variovorax* strains appear to belong to a clade of *Variovorax* strains that separates from the main bulk of strains, including most of the type strains (4). The ability to degrade and/or grow on linuron is unique for those strains within the *Variovorax* genus, indicating that they must have genetically adapted by acquiring the catabolic genes by horizontal gene transfer (HGT). This is supported by the observation that in strains SRS16 and WDL1, the catabolic genes are physically-linked with mobile genetic elements (MGE). In SRS16, the DCA catabolic genes are bordered by multiple insertion sequence (IS) elements^2^. The same applies to *hylA*, the *dca* cluster and the *ccd* cluster in strain WDL1^9^. Moreover, the three catabolic gene clusters in *Variovorax* sp. WDL1 reside on a large extra-chromosomal element that shows several plasmid features including gene functions for conjugation (8). However, how the constellation and the genetic context of the catabolic genes and their linkage with MGEs varies between different linuron-degrading *Variovorax* strains and how this relates to the geographic origin of the strains is yet unknown. Such knowledge will provide insight in the mechanisms that govern the functional evolution of genomes and especially those of organic xenobiotic degraders and more specifically of the genus *Variovorax*. This organism inhabits a wide variety of environments suggesting that it is prone to adaptation to new environmental constraints. To this end, we sequenced the complete genomes of six different linuron-degrading *Variovorax* strains isolated from distantly located geographical areas. We (i) re-analyzed the phylogenetic relationship between the strains and their phylogenetic position within the *Variovorax* genus, (ii) examined how their genomes differ with those of non-linuron degrading *Variovorax* strains emphasizing on the occurrence and types of MGEs and (iii) and compared the genetic constellation and context of the gene clusters involved in linuron metabolism to reveal how these traits were acquired among different strains.

## Results and discussion

### General genome features of linuron-degrading *Variovorax* strains

The full genome sequences of six linuron-degrading *Variovorax* sp. strains, i.e., WDL1 (5), SRS16 (2), PBL-H6, PBL-E5, PBS-H4 (4) and RA8 (3) were obtained. Their general genomic features are listed in Table 1. Strain WDL1 was recently found to consist of two subpopulations that only deviate in the presence of linuron or DCA degradation genes (9). In this study, the genome of one of these two subpopulations, i.e., the one carrying the *hylA* gene cluster was re-sequenced. The new sequence deviated slightly from the one reported by Albers et al. (9) The 5400 kb and 1240 kb replicons reported in (9) formed one chromosome of 6.7 Mbp while the 1380 kbp plasmid-like extrachromosomal replicon consisted of two replicons, i.e., pWDL1-1 (800 kbp) carrying linuron catabolic genes and pWDL1-2 (540 kbp). Chromosome sizes (ranging from 5.99 to 8.36 Mbp) and GC content (ranging from 66.24 to 66.86%) of the linuron-degrading strains were comparable to those of other *Variovorax* genomes. Sizes of other reported non-linuron degrading *Variovorax* genomes range between 4.31 to 9.24 Mbp (median: 7.2 Mbp) with GC contents of 64.6 to 69.6 (median: 67.4) (Table S1). All linuron-degrading *Variovorax* strains contained a relatively high number of extra-chromosomal elements (two to six), including smaller obvious plasmid replicons (20 to 70 kbp) but also larger replicons of more than 500 kbp). The GC content and codon usage of most of those larger extra-chromosomal elements substantially differed from those of the chromosome (Figure S1). They also did not contain any essential genes for cell viability, categorizing them rather as plasmids than as a second chromosome or chromid (10, 11). pPBL-E5-2 and pSRS16-3 showed GC-contents similar to those of the chromosome but did not contain genes for cell viability and carried plasmid-like replication modules, suggesting they are also plasmids. In contrast to the linuron-degrading *Variovorax* genomes, none of the non-degrading ones *Variovorax* strains for which genome sequences are available, carried plasmids. IncP-1 plasmids were though reported in three not phylogenetically-classified *Variovorax* isolates with unknown genome sequences. pHB44(12) and pBS64(12) were identified in *Variovorax* strains associated with the mycorrhizal fungus *Laccaria proxima*, and carry genes that increase the *Variovorax* host fitness by enabling metal ion transport and bacitracin resistance(13). pDB1(14) in *Variovorax sp*. DB1, carries genes for the biodegradation of the herbicide 2,4-dichlorophenoxyacetic acid (2,4-D). Another feature that distinguishes the linuron-degrading strains from the non-degraders is the occurrence of a high number of IS*1071* elements varying from four to seven copies in the degraders, while non-degraders did not carry any IS*1071*. IS*1071* is an insertion element that was first described bordering the 3-chlorobenzoate catabolic genes of *Pseudomonas* sp. BRC60 plasmid pBRC60(15). Since then it has been frequently associated with primarily catabolic genes in various organisms, especially β-proteobacteria (16). It often flanks the catabolic genes at both sites, forming a putative composite transposon which has been shown to translocate as a whole (15). The element has been suggested to play a primarily role in the acquisition and subsequent distribution of adaptive genes and especially catabolic functions in bacteria (16–18).

**Table 1:**
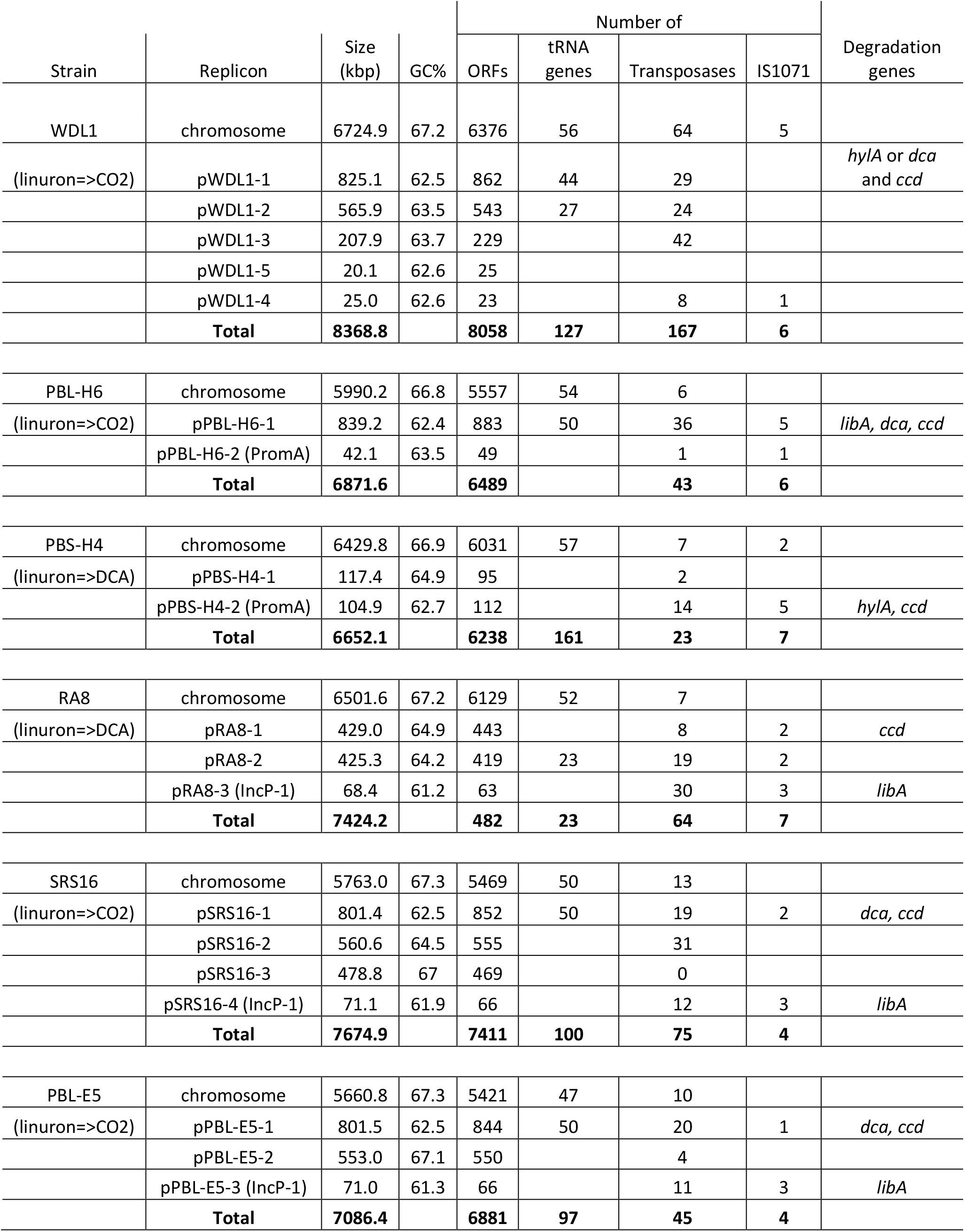
General properties of the newly sequenced *Variovorax* genomes

### Phylogenetic analysis of linuron-degrading *Variovorax* strains

The phylogenetic relatedness between the linuron-degrading *Variovorax* strains was determined by digital DNA:DNA hybridization values (dDDH) (Table S2). With the exception of PBL-E5 and SRS16, which represent the same species (dDDH value of 86%), all linuron-degrading *Variovorax* were designated as distinct species, and none of them belonged to any type species. Their phylogenetic divergence strongly indicates that the five degraders acquired linuron degradation genes independently as opposed to being derived from one common ancestral linuron degrader. Whole genome-based phylogeny (showed that the *Variovorax* species sequenced to date separated into two clades (clade 1 and clade 2), and that the linuron-degraders are very closely related species, all belonging to clade 1. The tree topology remained the same when only the linuron-degrader chromosomes were used (Figure S2). This separation largely replicated the 16S rRNA gene sequence-based phylogeny (Figure S3) with the exception of *V. soli* and *V. sp*. OV329. The low dDDH values of the *V. soli* genome with the degrader genomes (24.7-25.7%) however confirmed that these are distantly related. In addition to the linuron-degrading *Variovorax* strains, clade 1 included various other isolates but no type species. From those, a closed genome sequence was only available for the lignin-degrading soil isolate strain HW608 (19). Clustering of the linuron degrading strains was independent of either the geographical origin, the capacity to degrade linuron completely or partially to 3, 4-DCA, or the presence of specific catabolic genes involved in linuron biodegradation. Clade 2 contained the majority of the *Variovorax* strains, including the species *V. boronicumulans, V. soli, V. paradoxus, V. gossypi* and *V. guangxiensis*. Interestingly, in contrast to clade 2, non-linuron degrading strains from clade 1 were often associated with the catabolism of natural and anthropogenic organic compounds such as *Variovorax* sp. WS11 (an isoprene-degrading phyllosphere isolate (20)), KK3 (a 2,4-D-degrading freshwater isolate (21)), and JJ 1663 (an *N*-nitroglycine-degrading activated sludge isolate (22)). As such, including the linuron degraders, eight of the fourteen clade 1 strains were degraders of anthropogenic compounds. Clade 2, however, included only one xenobiotic-degrading isolate (one in 55 strains), i.e., *V. boronicumulans* J1(23) that degrades the neonicotinoid thiacloprid but only co-metabolically (24). These results indicate that the linuron-degrading strains belong to a *Variovorax* clade or originates from common ancestor that is/was more prone to genetic adaptation and hence specialization towards the biodegradation of anthropogenic compounds. The clade separation of strains with and without biodegradation capacity has not been observed before in other genera (25, 26).

### Plasmids hosted by linuron-degrading *Variovorax sp*

Phylogenetic analysis of the entire plasmid sequences clustered some of the plasmids identified in the linuron-degrading *Variovorax* sp. strains, into well-known but also novel plasmid groups (Figure 2). Some of the plasmids occur in multiple strains in which they are highly conserved (Figure S2). Table S3 shows overview of the replication and conjugation systems encoded on these plasmids. Twelve plasmids have type IV secretion system genes (T4SS) which facilitate conjugative transfer, but the origin of transfer (*oriT*) could not always be determined.

**Figure 1:**
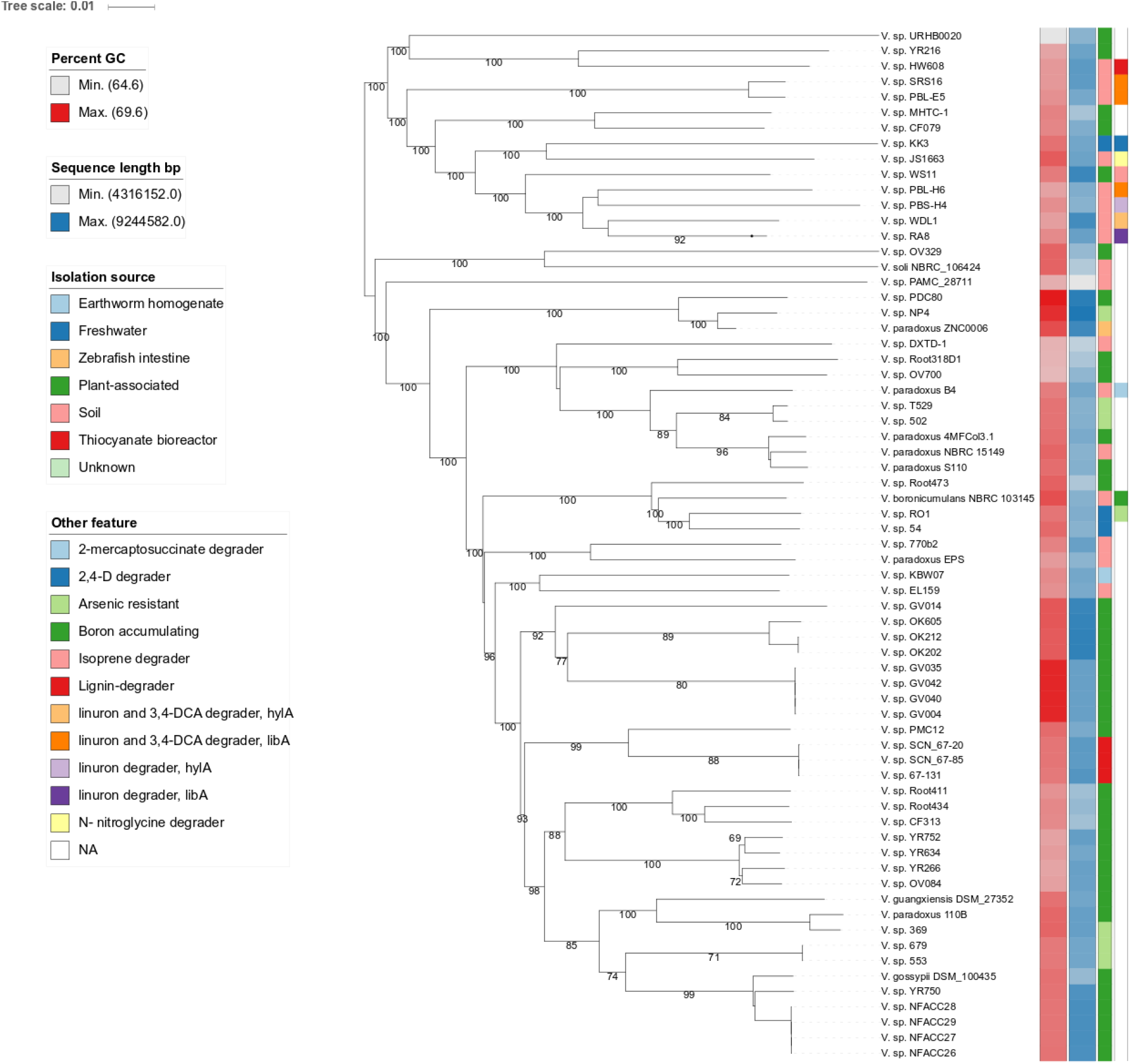
GBDP phylogenomic analysis of the *Variovorax* whole genome dataset. The branch lengths are scaled in terms of GBDP distance formula *d_5_*. The numbers above branches are GBDP pseudo-bootstrap support values from 100 replications, with an average branch support of 80.6%. Leaf labels are further annotated by their genomic G+C content, genome sequence length, phenotypic attributes and origin of isolation.

**Figure 2:**
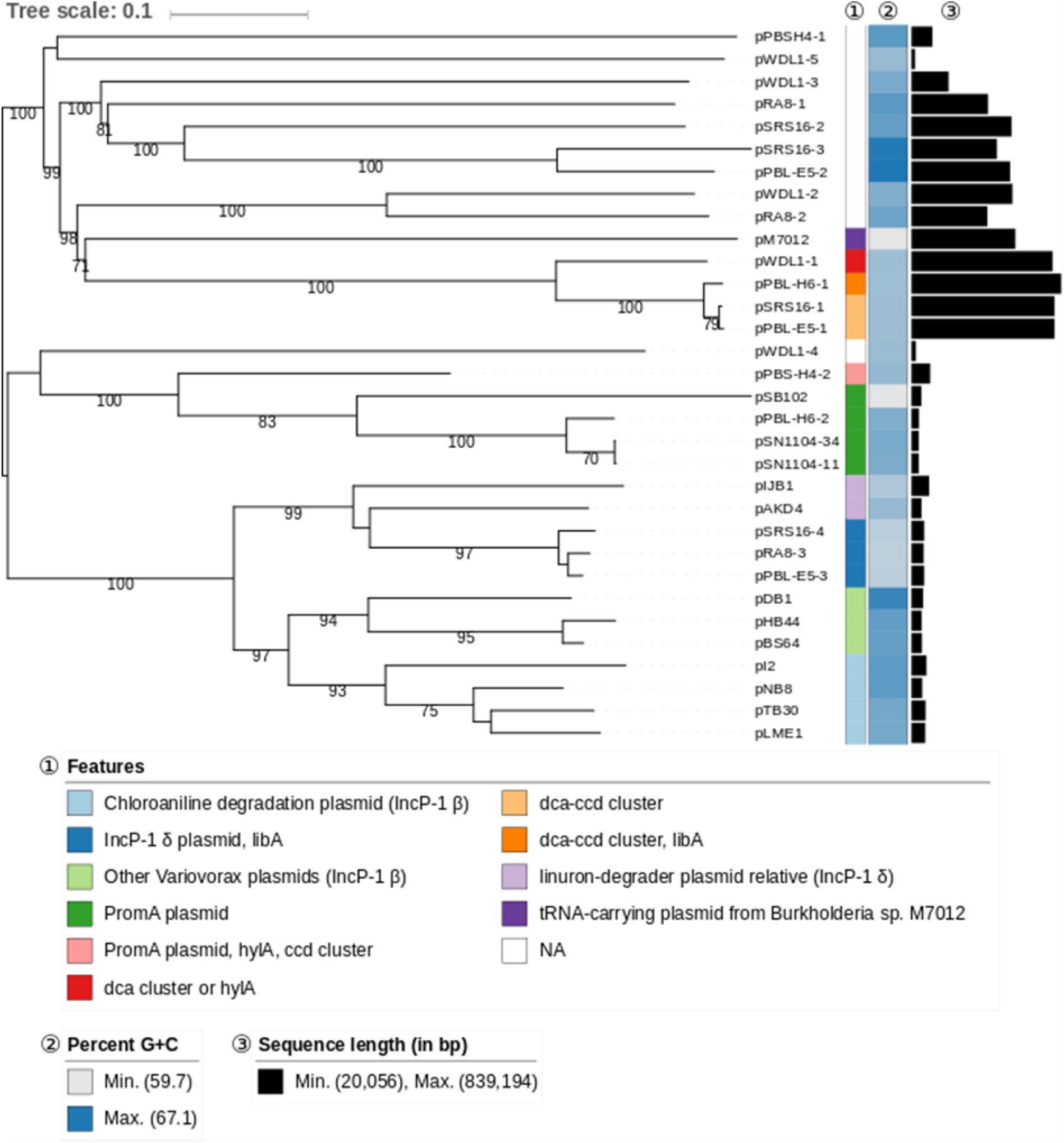
GBDP phylogenomic analysis of *Variovorax* plasmids and relevant relatives. The branch lengths are scaled in terms of GBDP distance formula *d6*. The numbers above branches are GBDP pseudo-bootstrap support values from 100 replications, with an average branch support of 91.3%. Leaf labels are further annotated by their genomic G+C content, length as well as special attributes. Previously-sequenced plasmids included in the study and their NCBI accession numbers are: pM7012 (NC_022995.1), pSB102 (AJ304453.1), pSN1104-34 (AP018708.1), pSN1104-11 (AP018707.1), pIJB1 (JX847411.1), pAKD4 (GQ983559.1), pDB1 (JQ436721.1), pHB44 (KU356988.1), pBS64 (KU356987.1), pL2 (JF274989.1), pNB8 (NC_019264.1), pLME1 (NC_019263.1) and pTB30 (NC_016968.1).

#### Known conjugative plasmids and their role in linuron degradation

Plasmids pPBL-E5-3, pRA8-3, and pSRS16-4 were classified as IncP-1δ plasmids. The three plasmids were 99% identical at the nucleotide (nt) level over the entire plasmid sequence and carry the *libA* locus between *trfA* and *oriV*, a known insertion hot spot for accessory genes in IncP-1 plasmids (16). Catabolic IncP-1δ plasmids have been reported before either isolated by means of exogenous isolation (27) or from isolates^17,^(29) all carrying 2,4-D degradation genes, between *trfA* and *oriV*. The above mentioned *Variovorax* plasmids pDB1(14), pHB44(12) and pBS64(12) are also IncP-1 plasmids, but belong to the IncP-1 β group.

Instead of IncP-1 plasmids, PBL-H6 and PBS-H4 carried self-transmissable broad host range PromA plasmids. pPBS-H4-2 carries *hylA* and the *ccd* gene cluster while pPBL-H6-2 carries an isolated IS*1071* transposase copy with inverted repeats (IR). pPBL-H6-2 and pPBS-H4-2 are most closely related to the PromA γ plasmids pSN1104-11 and pSN1104-34 (30) isolated by exogenous isolation, and hence pPBS-H4-2 and pPBL-H6-2 represent the first PromA γ plasmids obtained from isolates. Their presence in *Variovorax* extends the host range of PromA γ plasmids, as for PromA α and PromA β plasmids, to β-proteobacteria. pPBS-H4-2 is the first catabolic PromA plasmid, and is one of the few non-cryptic PromA plasmids(31). The often cryptic character of PromA plasmids has been the subject of a debate since it might harness their stability as they do not benefit the host fitness. It was suggested that they mainly support the conjugative transfer of other mobilizable replicons (31). The finding that PromA plasmids can carry catabolic genes shows that they, as is the case for IncP-1 plasmids, can indeed acquire and distribute genes beneficial for the host. The location of cargo genes in both pPBS-H4-2 and pPBL-H6-2 (near *virD2*) differs from this in other PromA plasmids (near *parA*)(32) and identifies the *virD* locus as an alternative hot spot for insertion of accessory genes in PromA plasmids.

#### Novel putatively conjugative plasmids in linuron-degrading Variovorax strains

Other plasmids than IncP1 and PromA plasmids were identified that carry homologues of TS44 genes (Table S3), i.e., pWDL1-3 and pWDL1-5, pRA8-1 and pSRS16-2. None of those plasmids categorized into a known plasmid group. Although these plasmids carried T4SS, the origin of transfer (oriT) and the type IV coupling protein (T4CP) could not always be identified, and a relaxase, which is necessary for conjugative transfer (33), could only be identified in pWDL1-3. pWDL1-3 carries a remarkably high number of 41 putative transposases albeit without IS*1071*, and several gene clusters for xenobiotic degradation. Among these is a gene cluster that encodes for homologues (40-43% identity) of proteins encoded by the *tphA1A2A3BR*–gene cluster for terephthalate degradation in *Comamonas sp*. E6(34) as well as for benzoate 1,2-dioxygenase subunits. pRA8-1 is distantly related to pWDL1-3 and carries the *ccdCFBD* operon. In addition, it contains homologues of genes for the biodegradation of non-chlorinated catechols, as well as cation efflux proteins CusABF (35) and cadmium transport protein CadA (36) flanked by an IS*1071* element. The small pWDL1-5 carries no cargo gene. A highly similar plasmid (99% nt identity, 72% coverage), also without cargo, is present in the chlorobenzene-degrading *Pandoraea pnomenusa* strain MCB032(37). The finding of this plasmid group in two different genera/families of the same bacterial order indicates its transferability within *Burkholderiales*. The Trb homologues encoded by pSRS16-2 as well as the *oriT* are highly similar (75-80%) to those encoded by IncP-1 plasmids. However, unlike IncP-1 plasmids, pSRS16-2 does not carry *trfA*, and its size (560 kbp) is much larger than IncP-1 plasmids. pSRS16-2 encodes for a broad range of functions, including 37 transport-related proteins, 18 proteins related to aromatic degradation and 31 transposases. We conclude that this plasmid represents a novel plasmid group, with conjugative transfer machinery similar to this of IncP-1 plasmids.

#### Non-transferrable plasmids pSRS16-3 and pPBL-E5-2

The closely related plasmids pPBL-E5-2 and pSRS16-3 carrying the *repB-parAB* replication system do not contain homologues of genes related to conjugal transfer, suggesting that they are not self-transmissable. These plasmids are different to the rest in that they have a GC content and codon usage similar to the chromosomes. Both carry distantly-related homologues of catabolic genes such as *tfdA* encoding conversion of 2,4-D (33% aa identity) in *Cupriavidus necator* JMP 134(38), and the *dmpKLMNOPQBCDEFGHI* gene cluster for phenol degradation (45-66% aa identity) in *Pseudomonas sp*. CF600) (39), as well as the *phnCDEGHIJKLMN* gene cluster for phosphonate uptake and degradation (45-62% aa identity) in *Eschericia coli* K12) (40).

#### t-RNA carrying megaplasmids of linuron-degrading *Variovorax sp*

Pairwise alignment showed that pPBL-H6-1, pSRS16-1, pPBL-E5-1 and pWDL1-1 are highly identical to one another. These show 99% nt identity over the entire sequence including the putative replication/partitioning module *repB*-*parAB* (Figure 3A). pPBL-H6-1 carries all three gene clusters required to convert linuron to 3-oxoadipate, while pSRS16-1 and pPBL-E5-1 only carry the *dcaQTA1A2BR* and *ccdCFDE* gene clusters. The pWDL1-1 variant sequenced in this study carries the *ccdCFDE* genes and the *hylA* gene. The proteins encoded by the *repB*-*parAB* module show only slight similarity to their nearest relatives (27, 48 and 39% aa similarity for RepB, ParA and ParB, respectively) and hence the four mega-plasmids might represent a new plasmid group, potentially specific for *Variovorax*. All four plasmids carry *traALBFHJDNUWG* and *trbCG* homologues, albeit with low similarity at aa level (30-43%) involved in conjugative transfer, suggesting that these may be transferrable plasmids. No putative relaxase was though found.

**Figure 3:**
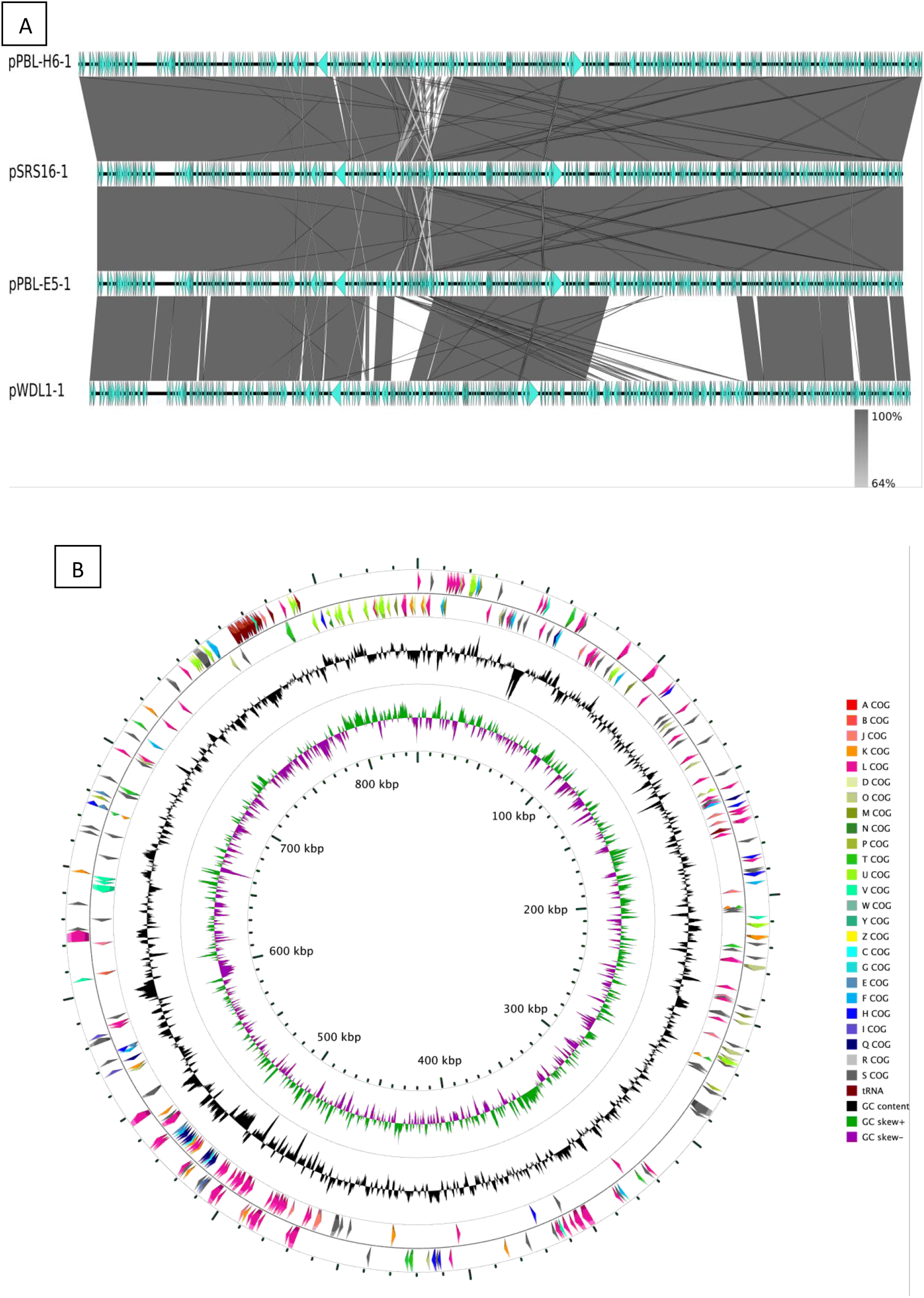
(A) Synteny and BLAST identity of the four Variovorax tRNA carrying megaplasmids. Shaded regions indicate the BLAST identity at the nucleotide level. (B) Circular representation of the pPBL-H6-1 as an example of a Variovorax tRNA carrying megaplasmid. The colored arrows represent the main COG categories that the proteins were assigned to, as well as the tRNA genes. G+C content and skew are represented in the inner circles.

Strikingly, the four megaplasmids carry tRNA genes that encode for all the proteinogenic aa codons. Unlike the scattered appearance of the tRNA genes located on the chromosome, the plasmid encoded tRNA genes are concentrated in one array (Figure 3B). Except for the tRNA encoding for codons glutamine (CAG) and arginine (CGC), all tRNAs on these plasmids are also present on the chromosome. In all four hosts, these two codons are preferred by both plasmids and the chromosome, however, multiple other tRNAs encode for these aa’s on the chromosome, suggesting that although the plasmid-encoded tRNAs may enhance gene expression they are not essential for expression of plasmid genes. pSRS16-1 further lacks the tRNA for valine (CAC), which is preferred by both the chromosome and plasmids; however this tRNA is present in pSRS16-2 as well as in the chromosome.

The presence of tRNA genes on large plasmids has been reported before in other bacteria (41–44)·. None of these plasmids however related to the tRNA-carrying *Variovorax* plasmids. In *Bifidobacterium breve* the tRNA-encoding plasmid improves gene expression from both the chromosome and the plasmid (43) while in *Anabaena sp*. PCC 7120 it is dispensable for growth (44). Other MGEs different from plasmids (45) as well as bacteriophages (46) encode for tRNA genes. In the acidophilic, bioleaching bacterium *Acidithiobacillus ferrooxidans*, the tRNA genes are located on an integrative-conjugative element and are likely not essential for growth (45).

Another feature that sets these plasmids apart is the presence of a CRISPR3-Cas cassette, which is identical (100% identity on nt level) in all of them. The cassette consists of a CRISPR array with 15 spacers, in addition to the genes encoding for a Cas6/Cse3/CasE-type endonuclease, the Cascade subunits CasA, CasB, CasC and CasD and the CRISPR-associated proteins Cas1 and Cas2, which is similar to the class I-E CRISPR-Cas systems (47). The CRISPR-defense system protects bacteria and archaea against MGEs and phages (48), and can be transferred horizontally (43, 49). Although the exact direct repeats of the CRISPR structure were also found in the *Serpentinomonas mccroryi* strain B1 genome (GCA_000828915.1), the spacer sequences did not have any match in the CRISPR databases. Interestingly, no CRISPRs were found in other *Variovorax* genomes available in public databases, with the exception of *Variovorax sp*. PDC80 (GCF_900115375.1), which carries a class I-F CRISPR-Cas system with spacers unrelated to those of the megaplasmids.

Other plasmids that carry tRNA genes in the linuron-degrading strains are plasmids pRA8-2 and pWDL1-2. These plasmids are distantly-related to pPBL-H6-1, pSRS16-1, pPBL-E5-1 and pWDL1-1 and also carry the tRNA genes in an array, however, their tRNAs do not encode for all proteinogenic amino acids and are all redundant. The two plasmids neither have functions for conjugal transfer nor carry catabolic genes but encode for putative heavy metal resistance, like the cobalt-zinc-cadmium efflux system encoded by *czcABCD* (50, 51), the copper-response two-component system encoded by *cusRS* and *cusABRS* (35), and mercury resistance encoded by *merACPTR* (52). Unlike pWDL1-2, pRA8-2 carries an IS*1071* element adjacent to a gene cluster encoding for homologues of the toxin-antitoxin system proteins DinJ-YafQ (53) and the antirestriction protein KlcA that play a role in ensuring plasmid stability.

### Genomic organization of linuron degradation genes among different *Variovorax* strains

We analyzed the presence, location and genomic context of *hylA* and *libA* genes for linuron hydrolysis, *dcaQTA1A2BR* genes for DCA conversion to 4,5-chlorocatechol(54) and *ccdCDEFR* genes for 4,5-dichlorocatechol degradation(1, 6) in each of the degraders genome. These genes were not present in any of the other publicly-available *Variovorax* genomes.

#### Genomic context of the linuron-hydrolysis genes *hylA* and *libA*

The linuron hydrolysis genes *hylA* and *libA* were highly conserved among the different strains. The six linuron degraders carried either *hylA* or *libA*, but never both. *hylA* is present in one copy in strains PBS-H4 and WDL1. As previously reported for WDL1, *hylA* in strain PBS-H4 makes part of a larger gene cluster of 13 ORFs that is highly conserved among the two hosting strains (99 to 100 % nt identity and complete synteny) and that is flanked at both sites by an IS*1071* composing a putative composite transposon (Figure 4A). The function of the *hylA* associated ORFs, and in particular the downstream ORFs, is currently unclear, but their conservative nature indicates that they play a role in linuron hydrolysis. Albers et al.(55) showed the upregulation of the downstream *luxRI* homologues when WDL1 is degrading linuron within a consortium. The *hylA* carrying composite transposon likely originated by inserting the *hylA* gene together with its downstream ORFs in a precursor composite transposon carrying the *iorAB* and *dca* gene clusters as suggested by the presence of *iorAB* and a *dcaQ* remnant directly upstream of *hylA*. Interestingly, the *dcaQ* gene that directly flanks *hylA*, is truncated at a different residue in WDL1 and in PBS-H4 (PBS-H4: nt position 749, WDL1: 689), which suggests that the *hylA* gene and its associated downstream ORFs were independently acquired by the composite transposons present in the two strains. However, this does not necessarily mean that IS*1071* recruited the *hylA* locus for WDL-1 and PBS-H4. The locus might have existed, though in different forms, before its recruitment by WDL1 and PBS-H4, as integrated into a composite transposon and might have been distributed as such.

**Figure 4:**
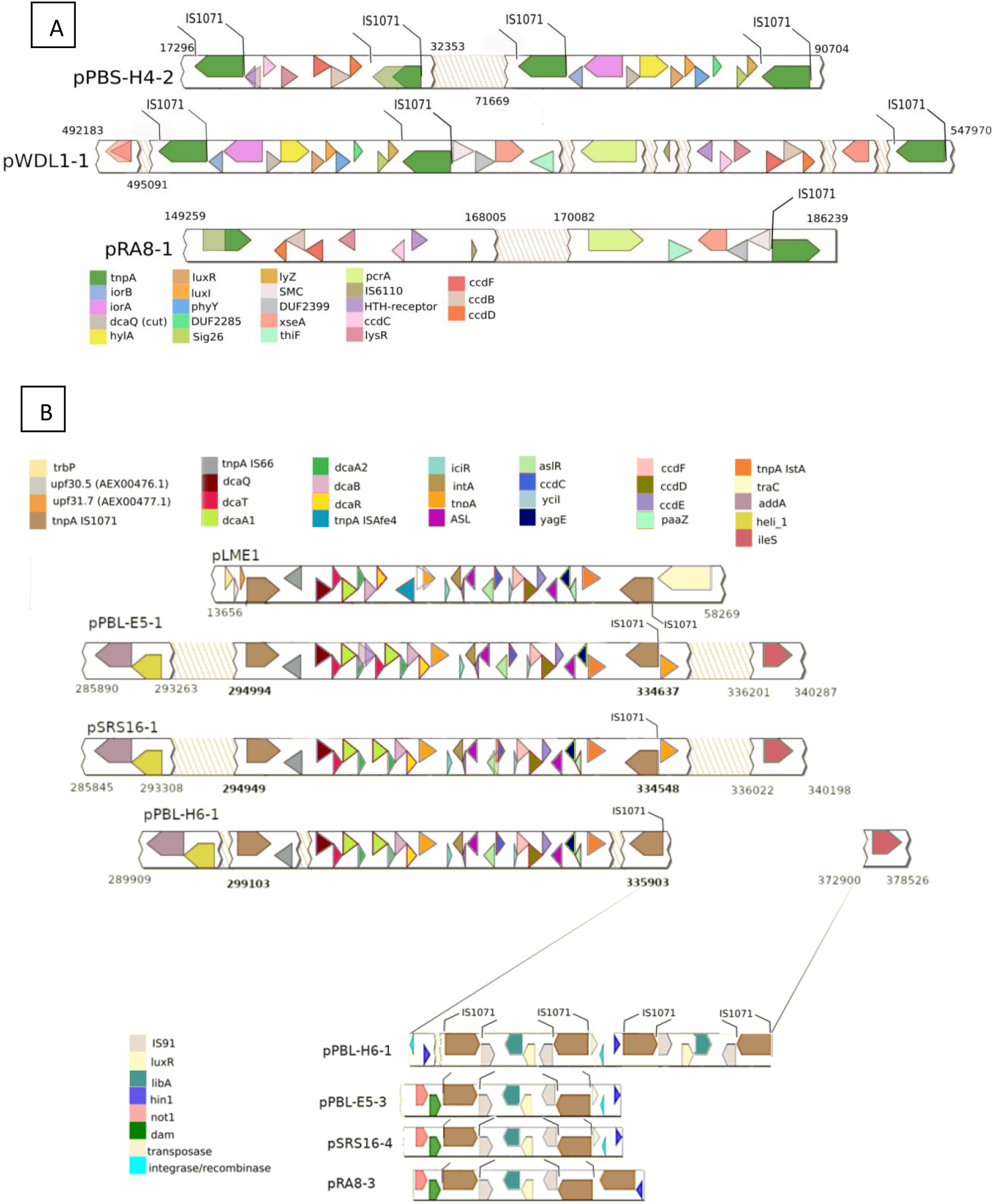
Catabolic clusters and their synteny. The genes with the same color code share 99-100% identity at the aa level. Shaded arrows indicate truncated genes, with the lighter-shade arrow indicating the full gene size. Broken likes indicate IS*1071* IRs. (A): Catabolic genes of WDL1, RA8 and PBS-H4. The *hylA* and *ccd* clusters on the plasmids pPBS-H4-2, pWDL1-1 and pRA8-1, as well as flanking genes are illustrated. Catabolic cluster locations are depicted for each plasmid in base pairs. (B) Catabolic genes of SRS16, PBL-H6 and PBL-E5. On the top panel, the *dca* and *ccd* clusters on plasmids pPBL-E5-1, pSRS16-1 and pPBL-H6-1 as well as the *Delftia acidovorans* plasmid pLME1 are illustrated, together with flanking genes and genes directly neighboring each IS*1071* element. The catabolic cluster locations and IS*1071* insertion positions are given for each plasmid in base pairs. In the lower panel, the *libA* gene-associated IS*1071* elements are illustrated for pPBL-H6-1, pPBL-E5-2, pSRS16-4 and pRA8-3. The insertion site of the *libA*-associated IS*1071* elements on pPBL-H6-1 is depicted with fading lines. For the other three plasmids, genes directly flanking the IS*1071* elements are given as well.

*libA* is present in SRS16, PBL-E5, RA8 and PBL-H6. This gene also makes part of a larger highly conserved gene cluster of four ORFs flanked at both borders by an IS*1071* element and hence composing a putative composite transposon structure (Figure 4B). In addition to *libA*, this gene cluster contains a *luxR*-family transcription regulator directly adjacent to *libA* followed by an IS*91*-family insertion element. The remarkable conservation of the *libA* locus including its integration into a composite transposon, might suggest that *Variovorax* recruited the *libA* locus rather through an already existing composite transposon structure carrying *libA*, as suggested above for the recruitment of the *hylA* locus, As reported above, the *libA* locus is carried by identical IncP-1 plasmids in strains PBL-E5, RA8 and SRS16. SRS16 and PBL-E5 were isolated from the same agricultural field, albeit at different time points, and hence this plasmid seems to play an important role in distributing the *libA* locus in that field. RA8 though was isolated in Japan, indicating that similar plasmids evolved at different locations, or that the BHR plasmid was transferred across a large geographic distance. In contrast, in PBL-H6, which originated from a Belgian agricultural field, the transposon is located on the t-RNA carrying megaplasmid pPBL-H6-1 as two flanking complete copies. As such, in PBL-H6, the *libA* composite transposon appears to be obtained by integrating in a replicon different from IncP-1 plasmids. However, PBL-H6 also harbors the PromA plasmid pPBL-H6-2 that carries a copy of the IS*1071* element. An ancestral version of the PromA plasmid might have been the initial carrier of the *libA* carrying composite transposon in PBL-H6. After the ancestral plasmid was recruited by PBL-H6, the transposon might have transposed to and duplicated in pPBL-H6-1, after which the *libA* gene cluster was lost from pPBL-H6-2 by homologous recombination of the IS*1071* element, leaving one copy behind. A similar phenomenon was previously shown for an IncP-1 plasmid carrying genes for atrazine biodegradation (56).

#### The *dca* and *ccd* clusters of linuron-degrading *Variovorax*

Similar to the *hylA* and *libA* genes, MGEs determine the genomic context of the *dca* and *ccd* genes. PBL-E5, PBL-H6 and SRS16 carry the entire *dca* and *ccd* clusters, which is consistent with their ability to degrade linuron and DCA. The *ccd* clusters of strains PBL-E5, PBL-H6 and SRS16 are on the 800 Mbp megaplasmids, directly downstream of the *dca* clusters (Figure 4B). In both strains, the entire *dca/ccd* gene cluster is bordered by IS*1071* at both sides, with two inward-directed IRs and one outward-directed IR missing. A similar composite transposon configuration including the *dca* and *ccd* genes is found in the chloroaniline degrader *Delftia acidovorans* LME1(54) (Figure 4B), except that PBL-H6 and SRS16 have two copies of the *dcaA1A2* genes and PBL-E5 two adjacent copies of the *dcaQTA1A2B* genes with one intact and one truncated *dcaB*. Other chloroaniline degraders like *Comamonas testosteroni* WDL7 and *Delftia acidovorans* B8c(54) carry a similar structure but lacking the *ccd* genes (Figure 4B). Amino acid-level similarity of the proteins encoded by *dcaQTA1A1BR* and *ccdCFDE* between the three *Variovorax* strains and LME1/WDL7/B8c is around 99% (Table S4). As for the *hylA* and *libA* loci, we hypothesize that the *dca/ccd* gene clusters were obtained by being already integrated into an ancestral composite transposon but that afterwards gene rearrangements occurred that explained the observed variations. . This is further supported by the observation that in the chloroanaline degrading *Comamonas/Delftia* strains, the composite transposon structures carrying *dca/ccd* genes are located on IncP-1 plasmids, while in the three *Variovorax* strains they are located on the 800 kbp t-RNA carrying megaplasmids. In case of PBL-H6, these are directly adjacent to the composite transposon structure carrying *libA* (see above). In all three strains the location of the composite transposons is the same, marking the corresponding location as a likely accessory gene insertion hot spot in the mega-plasmid.

The *dca* cluster and associated ORFs present on pWDL1-1 are also bordered by two IS*1071* elements, but the composite transposon does not contain a *ccd* cluster and the *dca* genes are rather related (99% nt identity) to the *tadQTA1A2B* encoding for conversion of aniline to catechol in *Delftia tsuruhatensis* AD9 (57). Also the *ccd* cluster, present on pWDL1-1, differs from those of PBL-E5, PBL-H6 and SRS16. This cluster is also bordered by IS*1071* elements and its location is in the direct vicinity of either *hylA* or the *dca* genes, depending on the WDL1 subpopulation (9). These genes are relatively distantly-related to other known catechol degradation genes (Table S4).

The *ccd* gene clusters in the non-DCA degraders PBS-H4 and RA8 are identical to those of WDL1. In PBS-H4, the *ccd* cluster is on the PromA plasmid pPBS-H4-2, that also bears the *hylA* carrying composite transposon. The *ccd* cluster is flanked by IS*1071* elements at both ends composing a putative composite transposon. Unlike pWDL1-1, where the putative transposons carrying the *hylA* and *ccd* cluster are directly adjacent to each other, on pPBS-H4-2 they are separated by other IS*1071* elements that don’t carry catabolic genes. In RA8, the *ccd* cluster is located on pRA8-1. This cluster is flanked by one truncated *tnpA*_IS*1071*_ (Figure 4A), without an IR, indicating that IS1071 also played a role in recruitment of the ccd cluster by pRA8-1. Overall, as for the *hylA*, *libA* and *dca* loci, the apparent strong conservation of the *ccd* genes in a composite transposon structure in different plasmid vehicles indicates again that these genes were recruited as a composite transposon structure that already contained the respective loci.

Overall, the analysis of the genetic context of the genes involved in linuron catabolism, either upstream or downstream functions in the pathway, indicates that IS*1071* insertion element play an essential role in the plasmid-associated acquisition of the catabolic genes and genetic adaptation of *Variovorax* toward the ability to degrade linuron. The phenomenon that IS*1071* elements are associated with genes involved in xenobiotic degradation was previously observed via both cultivation-dependent (7, 27, 58, 59) and cultivation-independent(16) methods. Subsequent inter- and intramolecular transposition of IS*1071* is thought to lead to the assembly of catabolic genes into a composite transposon structure with the recruited genes flanked by IS*1071* at both sites(60, 61). The new composite transposon structure can then move on itself between different replicons as shown for instance for Tn*5271*, the composite transposon consisting of the chlorobenzoate catabolic genes *cbaABC* flanked by IS*107*1, in *Alcaligines* sp BRC60(62). The strong conservative nature of the catabolic cargo in the putative composite transposon structures suggest that the recruitment of the different catabolic clusters in the linuron degrading *Variovorax* strains is rather due to the recruitment of already existing catabolic composite transposons rather than by the assembly process itself. Broad host range plasmids such as IncP-1 are known to distribute catabolic clusters in communities and often contain IS*1071* associated catabolic composite transposons (7, 54). Interestingly, each of the linuron degraders with the exception of WDL1, carries at least one plasmid of a well-known promiscuous plasmid group (IncP-1 or PromA). Their involvement in distributing linuron catabolic genes in the linuron degrading strains is suggested from the fact that these plasmids all carry at least one of the involved catabolic gene functions with the exception of pPBL-H6-2. However, as mentioned above, the latter carries an IS*1071* copy which might be obtained from internal recombination between two IS*1071* elements bordering a composite transposon as explained above. In addition, other plasmids of yet unknown type, carrying signs of conjugative features, were present. Likely, plasmids move around in a community, and pick up the composite transposons, for further transfer to their final hosts. As such, the BHR plasmids found in the linuron degrading strains might have, next to IS*1071*, functioned as a second crucial vehicle for acquisition of the catabolic genes. Interestingly, regarding catabolic functions, only a limited reservoir of catabolic loci was utilized for integration in the linuron catabolic pathways indicating that the choice of suitable catabolic functions for integration into a functional linuron catabolic pathway in *Variovorax* is limited, even on a worldwide scale. Curiously, these genes seem only to be recruited by a specific clade 1 of *Variovorax*, whose members, in addition to linuron, seem to be prone to genetic adaptation and hence specialization towards the biodegradation of other anthropogenic compounds. Apparently, for some reason, this clade is able to recruit and express foreign genes for xenobiotic biodegradation. Sequence analysis of other genomes within this clade, in addition to the linuron degraders, and comparison with the genome sequences of clade 2, might unravel the mechanisms involved in this special ability of catabolic gene recruitment.

## Materials and methods

### Genome sequencing, assembly and annotation

The details of the biomass and library preparation for genome sequencing are given in the supplementary text S1. Genome assembly was performed based on the PacBio reads by means of the RS_HGAP_Assembly.3 protocol included in SMRT Portal version 2.3.0 applying target genome sizes of 5 Mbp (PBL-E5), 15 Mbp (WDL1) and 10 Mbp (others). All assemblies showed one chromosomal contig, several extra-chromosomal contigs and several artificial contigs. Artificial contigs were removed from the assembly. The remaining contigs were circularized and assembly redundancies at the ends of the contigs were removed. ORFs on the replicons were ordered using *dnaA* (chromosome) or *repA*/*parA* (plasmids) as the first ORF. Error correction was performed by mapping the Illumina short reads onto finished genomes using bwa v. 0.6.2 in paired-end (sampe) mode using default settings(63) with subsequent variant and consensus calling using VarScan v. 2.3.6 (Parameters: mpileup2cns --min-coverage 10 --min-reads2 6 --min-avg-qual 20 --min-var-freq 0.8 --min-freq-for-hom 0.75 --p-value 0.01 --strand-filter 1 --variants 1 --output-vcf 1)(63). A consensus concordance of QV60 was reached. Automated genome annotation was performed using Prokka 1.8(64). Genomes were deposited at EMBL/ENA under the accession numbers LR594659-LR594661 (PBL-H6), LR594662-LR594665 (RA8), LR594666-LR594670 (SRS16), LR594671-LR594674 (PBL-E5), LR594675-LR594677 (PBS-H4), and LR594689-LR594694 (WDL1).

### Phylogenomic analysis of *Variovorax sp*. plasmids and genomes

The accession numbers of the genome and plasmid sequences used to construct the phylogenetic trees are listed in Table S1.

First, a phylogenomic analysis of the whole genome dataset was conducted at the nucleotide level using the truly whole-genome-based Genome-BLAST Distance Phylogeny method (GBDP)(65, 66)(67). Briefly, GBDP infers accurate intergenomic distances between pairs of genome sequences and subjects resulting distances matrices to a distance-based phylogenetic reconstruction under settings recommended for the comparison of prokaryotic genomes(65). The method is used by both the Genome-to-Genome Distance Calculator 2.1(65) and the Type Strain Genome Server(67).

A second phylogenetic analysis based on the plasmids’ amino acid sequences was conducted using GBDP as well, except that GBDP distance calculations were done under settings recommended for the analysis of bacteriophage sequences(68). The reason is that the sequence lengths of the plasmid sequences were in the same order of magnitude than bacteriophage sequences(68) and thus promised an equally good performance of the GBDP method when applied to plasmid data. The publicly-available plasmid sequences included in this study were selected based on their relatedness to the newly-sequenced plasmids, in order to allocate them into known plasmid groups.

Regarding both analyses, a balanced minimum evolution tree was inferred using FastME v2.1.4 with SPR postprocessing each(69). 100 replicate trees were reconstructed in the same way and branch support was subsequently mapped onto the respective tree(66).

For the 16S rRNA gene sequence-based phylogeny, the whole 16S rRNA gene sequences were retrieved from the SILVA database(70), and aligned with the SINA aligner(70). The phylogenies were inferred on the GGDC web server(65) using the DSMZ phylogenomics pipeline (https://ggdc.dsmz.de/phylogeny-service.php). Maximum likelihood (ML) and maximum parsimony (MP) trees were inferred from the alignment with RAxML(70) and TNT(70), respectively. For ML, rapid bootstrapping in conjunction with the autoMRE bootstopping criterion(70) and subsequent search for the best tree was used; for MP, 1000 bootstrapping replicates were used in conjunction with tree-bisection-and-reconnection branch swapping and ten random sequence addition replicates. The sequences were checked for a compositional bias using the X^2^ test as implemented in PAUP*(70). All phylogenetic trees were visualized with iTOL(71).

### Analysis of *Variovorax sp*. plasmids

The codon usage of chromosomes and plasmids was calculated with CompareM v. 0.0.23(72). Subsequently, a PCA was conducted and the principle components were hierarchically clustered using the Ward’s criterion with FactoMineR FactoMineR v. 1.36(73). The replication origins as well as the type IV secretion systems were predicted with oriTfinder v. 1.1(70).

The COG categories on pPBL-H6-1 was determined with eggNOG(74). The circular representation was drawn with CGview(75) (Figure 3A), and the BLAST-based comparative analysis illustration was generated with Easyfig(76) (Figure 3B). SimpleSynteny(77) was used to determine the positions of the catabolic clusters and associated ORFs, and draw the catabolic cluster illustrations.

## Acknowledgements

We thank Sebastian S. Sørensen and Koji Satsuma for the donation of strains SRS16 and RA8, Simone Severitt for technical assistance, and Markus Göker, Jörn Petersen, Kornelia Smalla and Isabel Schober for valuable discussions regarding the manuscript. JMK was supported by Deutsche Forschungsgemeinschaft within “Sonderforschungsbereich TRR 51”. The authors acknowledge the use of de. NBI cloud and the support by the High Performance and Cloud Computing Group at the Zentrum für Datenverarbeitung of the University of Tübingen and the Federal Ministry of Education and Research (BMBF) through grant no 031 A535A.

## Supplementary information

Supplementary Text S1: Library preparation for PacBio and Illumina sequencing:

Biomass for genome sequencing of the six linuron-degrading Variovorax was obtained from 10 mL cultures grown in R2B medium supplemented with 10 mg/L linuron (Sigma-Aldrich, analytical standard) at 20 °C till an OD600 of 0.8-1. The linuron and/or DCA degradation phenotype of the cultures was assessed by monitoring the compounds concentration as previously described(1). DNA was isolated from the cultures using Qiagen Genomic-tip 100/G (Qiagen, Hilden Germany) kit according to the manufacturer’s instructions. For long read sequencing, 15 kbp libraries were prepared according to the SMRTbell™ template preparation protocol of PacificBiosciences (Menlo Park, USA), following the Procedure & Checklist – Greater Than 10 kbp Template Preparation. Briefly, 8 μg genomic DNA was sheared using g-tubes™ from Covaris (Woburn, USA). DNA was end-repaired and ligated overnight to hairpin adapters using the DNA/Polymerase Binding Kit P6 (Pacific Biosciences, Menlo Park, USA). BluePippin™ Size-Selection gel cassettes were used to select for DNA fragments greater than 4 kbp according to the manufacturer’s instructions (Sage Science, Beverly, MA, USA). Conditions for annealing of sequencing primers and binding of polymerase to purified SMRTbell™ template were assessed with the Calculator in RS Remote (PacificBiosciences, Menlo Park, USA). One SMRT cell for sequencing was used per strain on a PacBio RSII apparatus (PacificBiosciences, Menlo Park, USA) taking one 240-minutes movie with exception of strain RA8, for which three SMRT Cells were used. For short read sequencing, short insert libraries were created using the Illumina Nextera XT DNA Library Prep Kit (Illumina, San Diego, USA) and pair-end (2 X 151 bp) sequenced on an Illumina NextSeq 550 (Illumina, San Diego, USA).

**Table S1:**
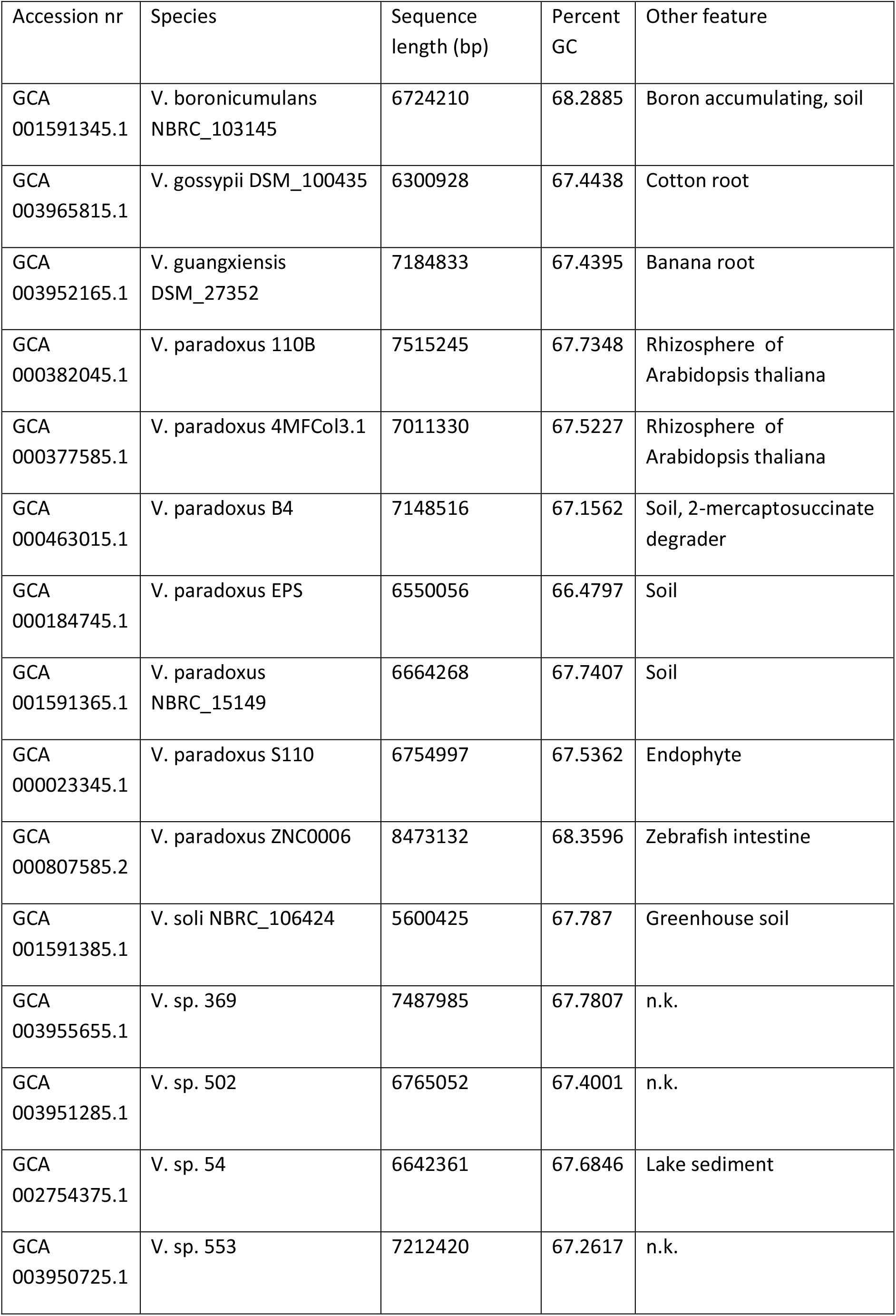
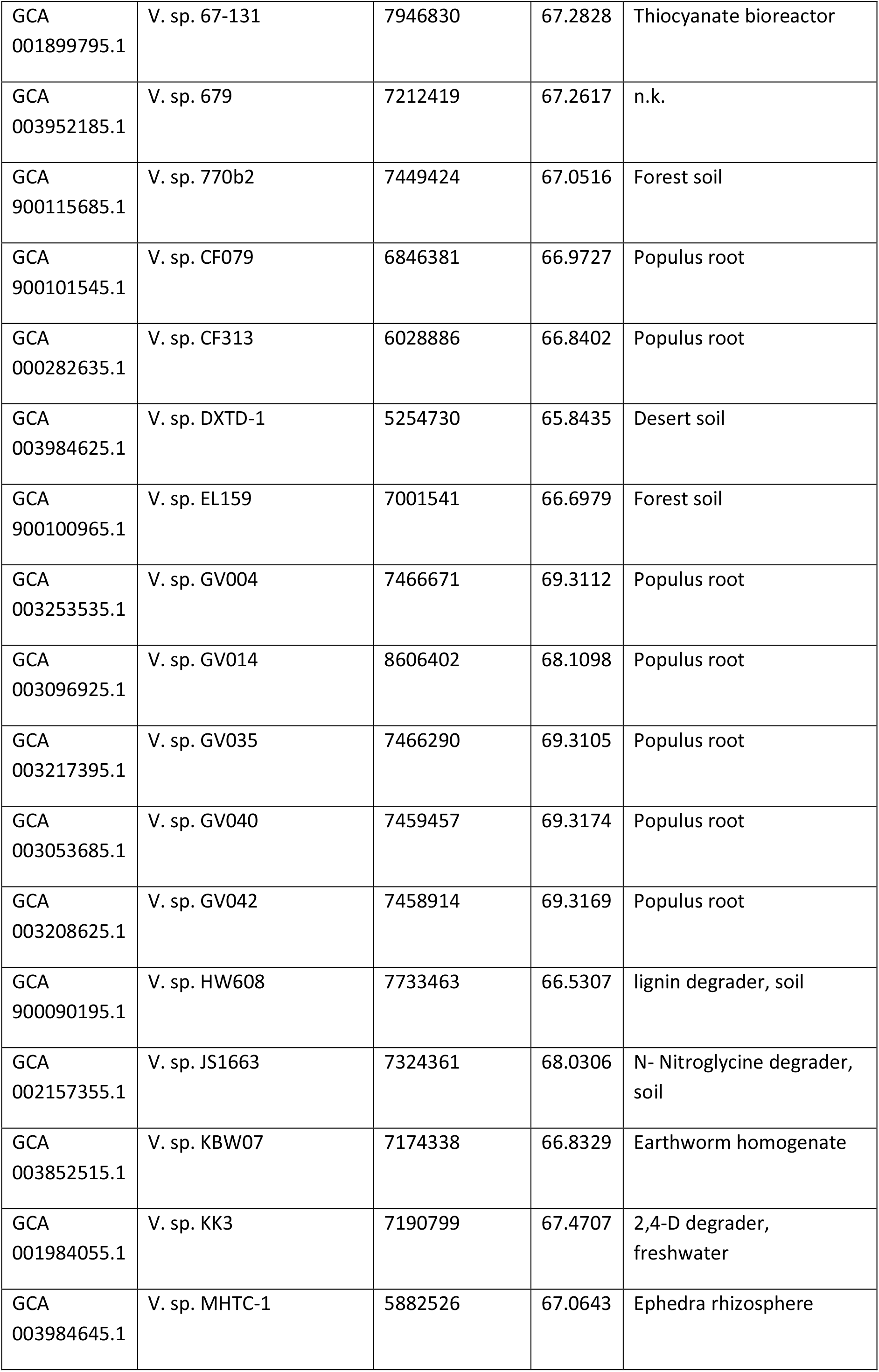
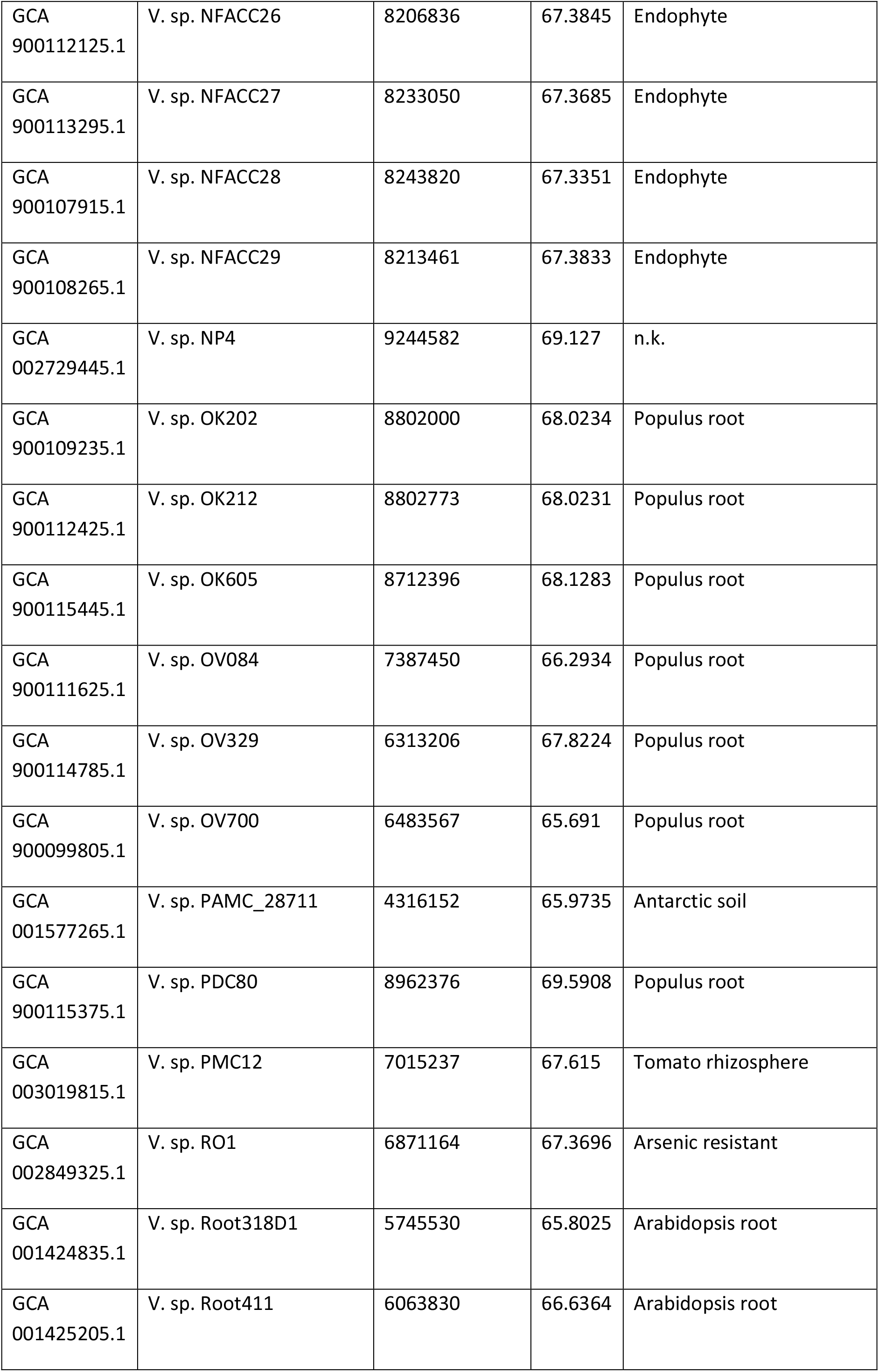
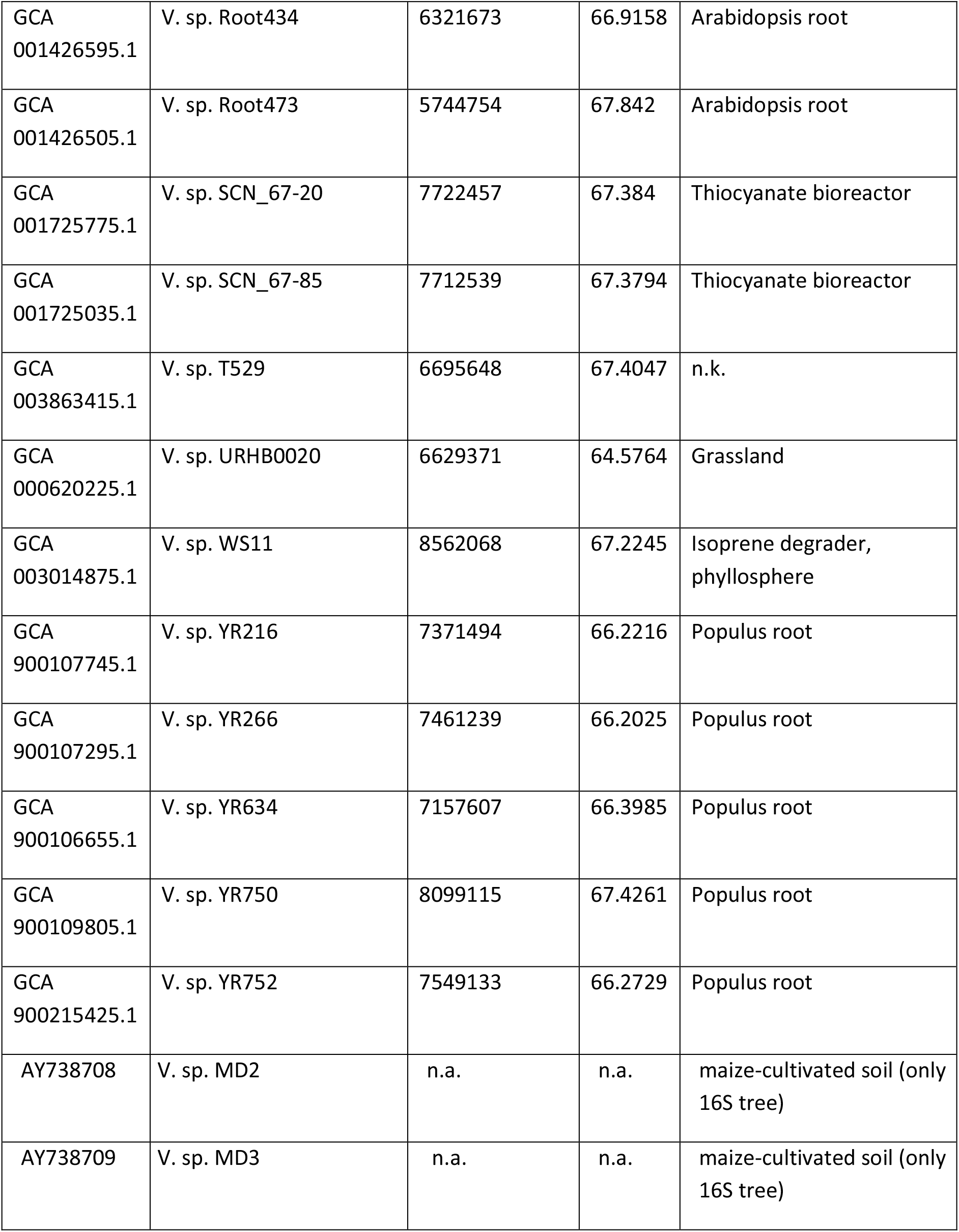
*Variovorax* genomes used in this study.

**Table S2:** Digital DNA-DNA hybridization values calculated by the type strain genome server (tygs.dsmz.de)(1). A value of 70% of higher indicates that the two microorganisms belong to the same species(2). C.I.: confidence interval.

| Query strain | Subject strain | dDDH (%) | C.I. (%) |
| --- | --- | --- | --- |
| PBL-E5 | SRS16 | 86 | [83.3 - 88.3] |
| RA8 | WDL-1 | 51.2 | [48.6 - 53.9] |
| PBL-H6 | WDL-1 | 44.9 | [42.3 - 47.4] |
| PBL-H6 | RA8 | 43.1 | [40.6 - 45.7] |
| PBS-H4 | RA8 | 40.1 | [37.7 - 42.7] |
| PBL-H6 | PBS-H4 | 39.9 | [37.4 - 42.4] |
| PBS-H4 | WDL-1 | 39.3 | [36.8 - 41.8] |
| PBL-E5 | PBL-H6 | 32.2 | [29.8 - 34.7] |
| PBL-H6 | SRS16 | 32.1 | [29.7 - 34.6] |
| PBL-E5 | WDL-1 | 30.6 | [28.2 - 33.1] |
| SRS16 | WDL-1 | 30.4 | [28.0 - 32.9] |
| PBL-E5 | RA8 | 26.9 | [24.6 - 29.4] |
| RA8 | SRS16 | 26.8 | [24.5 - 29.3] |
| PBL-E5 | PBS-H4 | 25.3 | [23.0 - 27.8] |
| PBS-H4 | SRS16 | 25.3 | [22.9 - 27.8] |

1. Meier-Kolthoff JP, Göker M. 2019. TYGS is an automated high-throughput platform for state-of-the-art genome-based taxonomy. *Nat Commun* 10:2182.
2. Liu Y, Lai Q, Göker M, Meier-Kolthoff JP, Wang M, Sun Y, Wang L, Shao Z. 2015. Genomic insights into the taxonomic status of the *Bacillus cereus* group. *Sci Rep* 5:14082.

**Table S3:** Overview of plasmid replication and conjugation systems. T4SS= type IV secretion system, T4CP= type IV coupling protein, n.f.= not found. P=present. For the relaxase, T4CP and T4SS. The % identity on aa level to the nearest relative, as well as the accession number are given.

| Plasmid | Relaxase | T4CP | T4SS | ori<br>T | Replication<br>genes |
| --- | --- | --- | --- | --- | --- |
| pPBL-E5-1<br>pWDL1-1<br>pSRS16-1<br>pPBL-H6-1 | n.f. | 34%<br>YP_001911165 | TraALBFHJDNUW,<br>TrbC, VirB4 | n.f. | RepB-ParAB |
| pSRS16-2 | TraI 49%<br>YP_195891 | 72%<br>NP_990928.1 | TrbLJIHGFEDCB,<br>TraJ, VirB1 | P | RepB-ParAB |
| pSRS16-3<br>pPBL-E5-2 | n.f. | n.f. | n.f. | n.f. | RepB-ParAB |
| pSRS16-4<br>pRA8-2<br>pPBL-E5-3 | TraI 100%<br>YP_006965894.1 | 100%<br>NP_990928.1 | TrbBCDEFGHIJKLN.<br>TraXF | P | IncP-1 |
| pPBL-H6-2<br>pPBS-H4-2 | TraS 79%<br>CAC79161 | 67%<br>YP_001672044 | VirB123456891011,<br>VirD4 | P | RepA |
| pRA8-1 | Rel 81%<br>SDZ72275.1 | 45%<br>WP_010895213 | VirB23456891011,<br>VirD4 | n.f. | RepB-ParAB |
| pRA8-2 | n.f. | n.f. | n.f. | n.f. | RepB-ParAB |
| pPBS-H4-1 | TraA 37%<br>AAV52093 | n.f. | n.f. | n.f. | RepA |
| pWDL1-2 | n.f. | n.f. | n.f. | n.f. | RepB-ParAB |
| pWDL1-3 | Rel 84%<br>WP_093180082.1 | 45%<br>WP_010895213 | VirB23456891011,<br>VirD4, TrbB | n.f. | RepAB |
| pWDL1-4 | n.f. | n.f. | n.f. | n.f. | ParAB |
| pWDL1-5 | n.f. | 38% AEY63616 | VirB568, TraD | n.f. | repA |
| chrWDL1 |  |  | TrbBCDEJLFGI,<br>VirD4, TraF |  |  |
| chrSRS16 |  |  | TrbBCDEJLFGI,<br>VirD4 |  |  |
| chrRA8 |  |  | n.f. |  |  |
| chrPBL-H6 |  |  | TrbBCDEJLFGI,<br>VirD4 |  |  |
| chrPBL-E5 |  |  | TrbBCDEJLFGI,<br>VirD4 |  |  |
| chrPBS-H4 |  |  | TrbBCDEJLFGI,<br>VirD4 |  |  |

**Table S4:** Presence of the catabolic proteins in each strain, their aa level identity to the nearest relative, and the accession number of the nearest relative.

|  | PBL-H6 | PBL-E5 | SRS16 | WDL1 | PBS-H4 | RA8 |
| --- | --- | --- | --- | --- | --- | --- |
| **HylA** | no | no | no | yes | yes | no |
| **LibA** | yes | yes | yes | no | no | yes |
| **DcaQ** | yes (100% identical to AEX00653.2 DcaQ [*Delftia acidovorans*]) | yes (see PBL-H6) | yes (see PBL-H6) | yes (99% identical to WP_047349924.1 DcaQ [*Delftia tsuruhatensis*]) | no | no |
| **DcaT** | yes (100% identical to AEX00654.1 DcaT [*Delftia acidovorans*]) | yes (see PBL-H6) | yes (see PBL-H6) | yes (98% identical to AAX47240.1 DcaT [*Delftia tsuruhatensis*]) | no | no |
| **DcaA1** | yes (100% identical to AEX00655.1 DcaA1 [*Delftia acidovorans*]) | yes (see PBL-H6) | yes (see PBL-H6) | yes (98% identical to AAX47241.1 DcaA1 [*Delftia tsuruhatensis*]) | no | no |
| **DcaA2** | yes (AEX00656.1 DcaA2 [*Delftia acidovorans*]) | yes (see PBL-H6) | yes (see PBL-H6) | yes (99% identical to AAX47242.1 DcaA2 [*Delftia tsuruhatensis*]) | no | no |
| **DcaB** | yes (AEX00657.1 DcaB [*Delftia acidovorans*]) | yes (see PBL-H6) | yes (see PBL-H6) | yes (99% identical to AAX47243.1 DcaB [*Delftia tsuruhatensis*]) | no | no |
| **DcaR** | yes (100% identical to AEX00658.1 DcaR [*Delftia acidovorans*]) | yes (see PBL-H6) | yes (see PBL-H6) | no | no | no |
| **CcdC** | yes (100% identical to WP_015060619.1 chlorocatechol 1,2-dioxygenase [*Delftia acidovorans*]) | yes (see PBL-H6) | yes (see PBL-H6) | yes (76% identical to WP_102776360.1 catechol 1,2-dioxygenase [*Achromobacter pulmonis*]) | yes (see WDL1) | yes (see WDL1) |
| **CcdF** | yes (100% identical to WP_015060622.1 maleylacetate reductase [*Delftia acidovorans*]) | yes (see PBL-H6) | yes (see PBL-H6) | yes (76% identical to AYE88726.1 chloromaleylacetate reductase [*Diaphorobacter sp.*]) | yes (see WDL1) | yes (see WDL1) |
| **CcdD** | yes (100% identical to WP_015060623.1 | yes (see PBL-H6) | yes (see PBL-H6) | yes (78% identical to AYE88727.1 | yes (see | yes (see |

|  | chloromuconate<br>cycloisomerase [*Delftia*<br>*acidovorans*]) |  |  | dichloromuconate<br>cycloisomerase<br>[*Diaphorobacter sp.*] ) | WDL1) | WDL1) |
| --- | --- | --- | --- | --- | --- | --- |
| **CcdE** | yes (100% identical to<br>WP_015060624.1<br>dienelactone hydrolase<br>[*Delftia acidovorans*] ) | yes (see<br>PBL-H6) | yes (see<br>PBL-H6) | yes (74% identical to<br>AYE88728.1<br>chlorodienelactone hydrolase<br>[*Diaphorobacter sp.*] ) | yes<br>(see<br>WDL1) | yes<br>(see<br>WDL1) |

**Figure S1:**
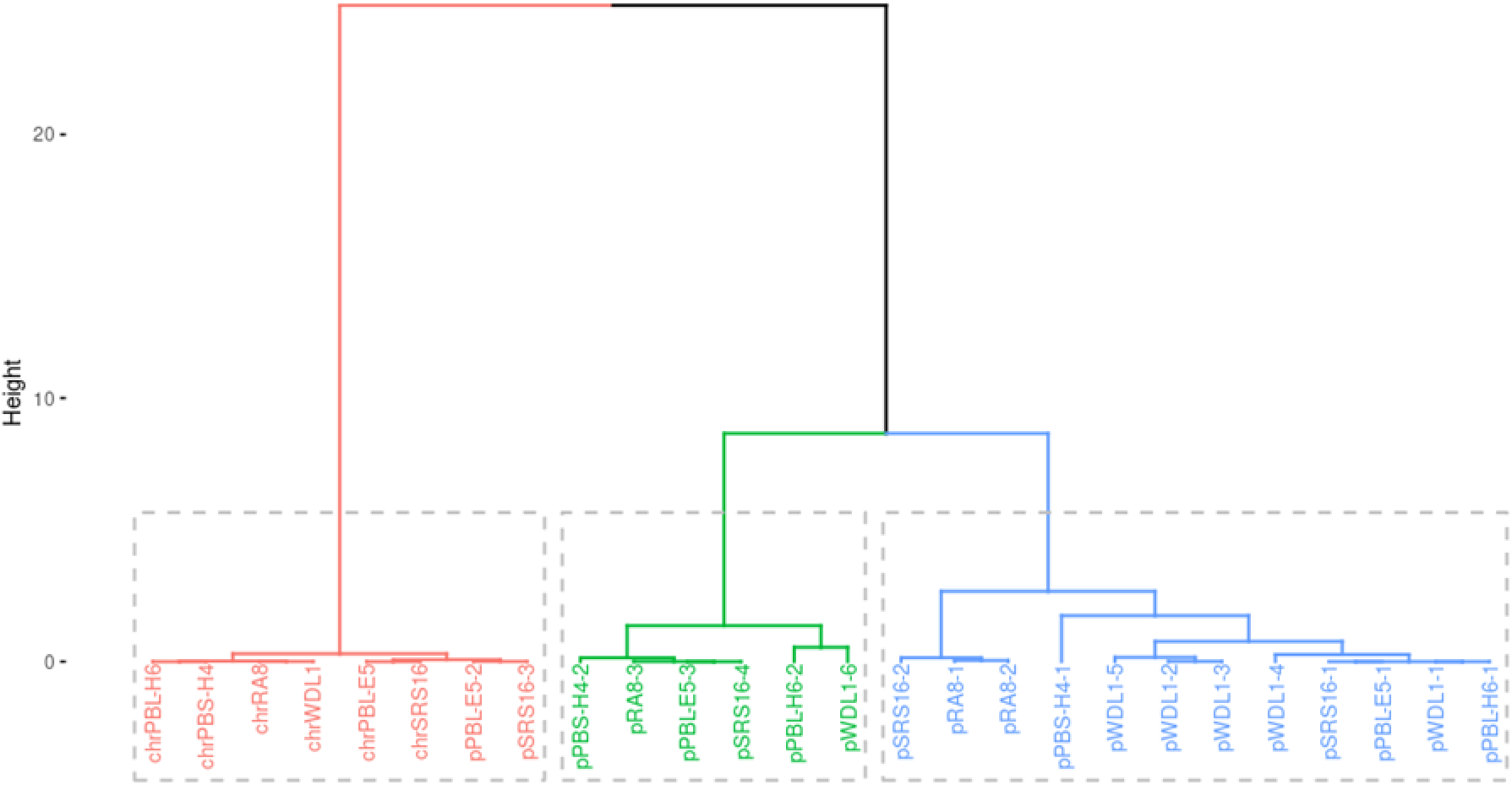
Hierarchical clustering of codon usage of all genomic elements

**Figure S2:**
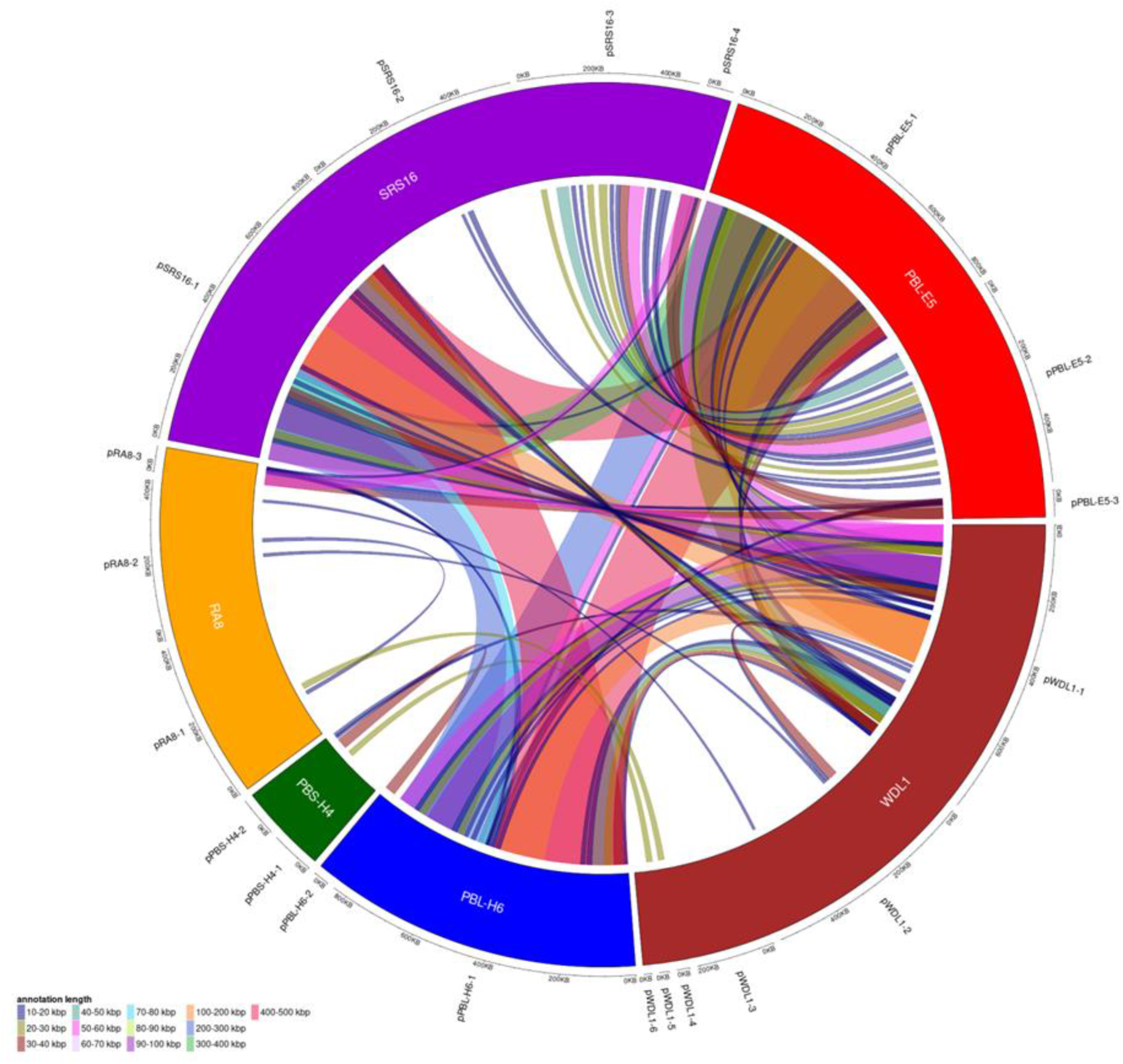
Circular representation of highly-similar regions between the plasmids of the degrading Variovorax genomes. On the ring, the different plasmids are shown, that are linked according to the length of the matching regions. The alignment was calculated with BlastN(2) (evalue < 1e-10, identity >= 95%), only alignment length of at least 10 kbp were taken into account. The circular plasmid comparison was generated with R v. 3.5.2 (https://www.R-project.org/) and the Bioconductor package circlize v. 0.4.5(3).

**Figure S3:**
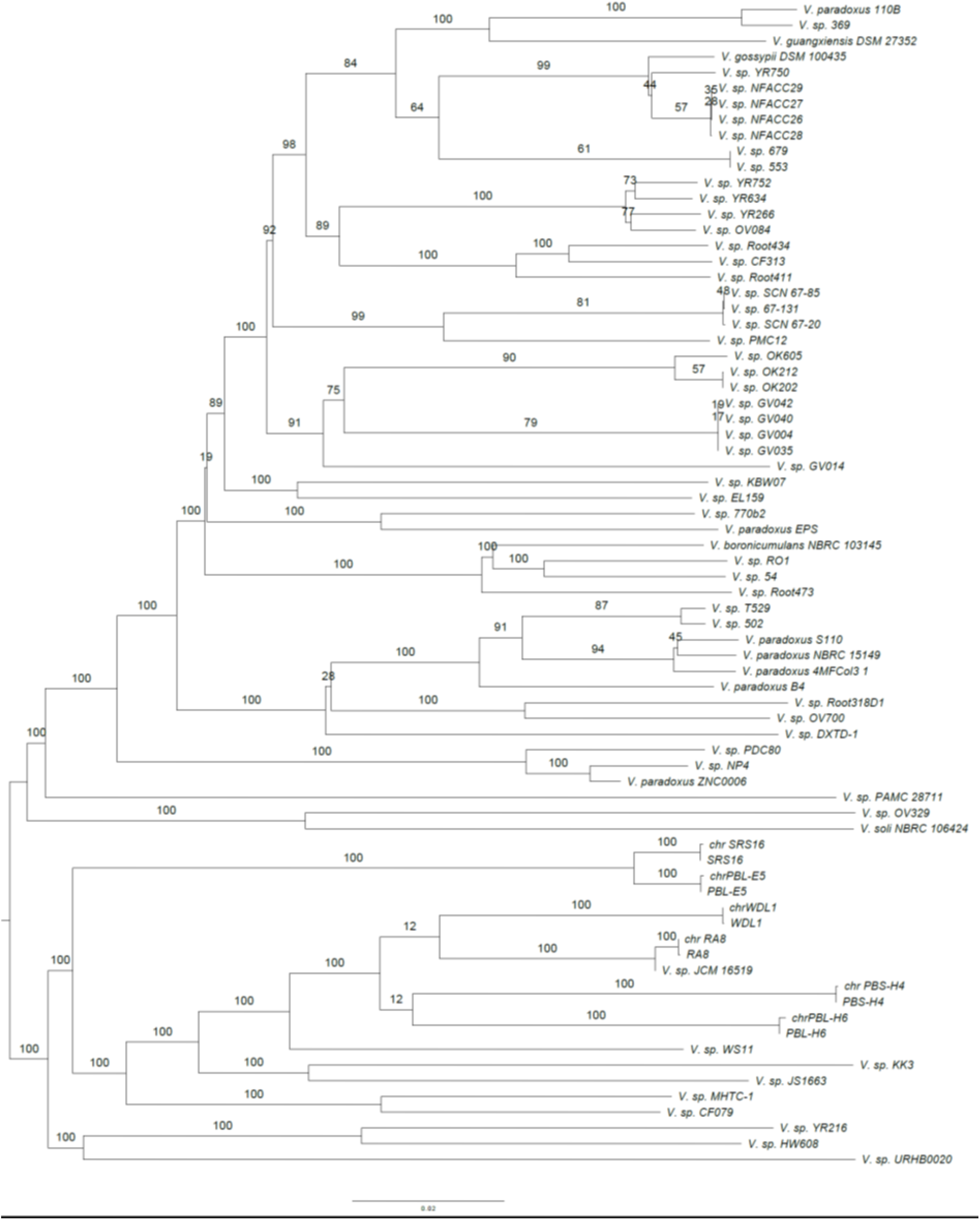
GBDP phylogenomic analysis of the Variovorax chromosome dataset. The branch lengths are scaled in terms of GBDP distance formula d5. The numbers above branches are GBDP pseudo-bootstrap support values from 100 replications, with an average branch support of 80.6%. For the linuron-degrading Variovorax, whole genomes were included in the calculation for comparison.

**Figure S4:**
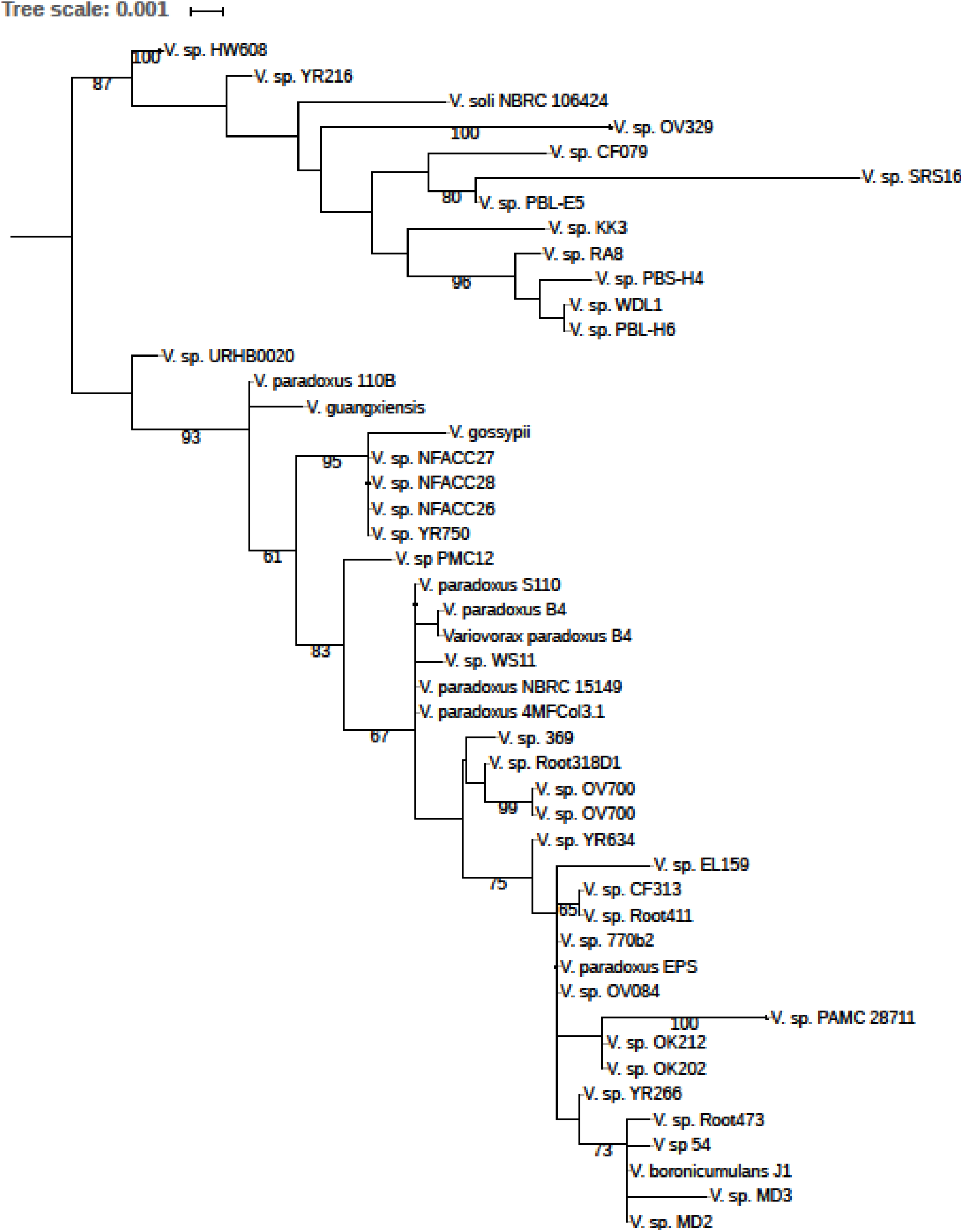
16S-based phylogeny of Variovorax. ML tree inferred under the GTR+CAT model and rooted by midpoint-rooting. The branches are scaled in terms of the expected number of substitutions per site. The numbers above the branches are support values when larger than 60% from ML bootstrapping.

